# BgTEP: an antiprotease involved in innate immune sensing in *Biomphalaria glabrata*

**DOI:** 10.1101/308130

**Authors:** Anaïs Portet, Richard Galinier, Silvain Pinaud, Julien Portela, Fanny Nowacki, Benjamin Gourbal, David Duval

**Author notes:** Equal contributions: AP & RG. corresponding author: David Duval, UMR5244 IHPE, Université de Perpignan, 58 avenue Paul Alduy, 66860 Perpignan.

## Abstract

Insect Thioester-containing protein (iTEP) is the most recently defined group among the TEP superfamily. TEPs are key components of the immune system, and iTEPs from flies and mosquitoes were shown to be major immune weapons. Initially characterised from insects, TEP genes homologous to iTEP were further described from several other invertebrates including arthropods, cniderians and mollusks albeit with few functional characterisations. In the freshwater snail *Biomphalaria glabrata*, a vector of the schistosomiasis disease, the presence of a TEP protein (BgTEP) was previously described in a well-defined immune complex involving snail lectins (FREP) and schistosome parasite mucins (SmPoMuc).

To investigate the potential role of BgTEP in the immune response of the snail, we first characterised its genomic organisation and its predicted protein structure. A phylogenetic analysis clustered BgTEP in a well-conserved subgroup of mollusk TEP. We then investigated the BgTEP expression profile in different snail tissues, and followed immune challenges using different kinds of intruders during infection kinetics. Results revealed that BgTEP is particularly expressed in hemocytes, the immune-specialised cells in invertebrates, and is secreted into the hemolymph. Transcriptomic results further evidenced an intruder-dependent differential expression pattern of BgTEP whilst interactome experiments showed that BgTEP is capable of binding to the surface of different microbes and parasite either in its full length form or in processed forms.

Through this work, we report the first characterisation of a snail TEP. Our study also reveals that BgTEP may display an unexpected functional dual-role. In addition to its previously characterised anti-protease activity, we demonstrate that BgTEP can bind to the intruder surface membrane, which supports a likely opsonin role.

## Introduction

Thioester-containing proteins (TEPs) are large secreted glycoproteins characterised by the presence of a unique intrachain β-cysteinyl-γ-glutamyl thioester bond (Williams and Baxter, 2014). In native TEPs this bond is unreactive, but following proteolytic activation, change in temperature or aqueous conditions, the thioester bond becomes reactive and can bind closely accessible hydroxyl or amine groups that are present at the surface of many biological entities including pathogens (Shokal and Eleftherianos, 2017).

The canonical intrachain thioester bond (GCGEQ) was originally described in the human alpha-2-macroglobulin (A_2_M), protease inhibitor and C3 protein, central components of the complement system (Dodds and Law, 1998). Since then, members of the TEP superfamily have been identified and characterised in numerous distant phyla from Ecdysozoans and Lophotrochozoans to Deuterostomes. The TEP superfamily is divided into two subfamilies displaying distinct functions, complement factors and A_2_M (Shokal and Eleftherianos, 2017), supported by the presence of anaphylatoxin (ANA) and C-terminal C345C domains only in members of the complement factor subfamily. The complement factors subfamily contains human C3, C4 and C5 proteins, and their orthologs. The A_2_M subfamily comprises A_2_M, pregnancy zone protein-like (PZP), complement C3 and PZP A_2_M domain-containing 8 (CPAMD8), and cell surface glycoprotein CD109. In addition, two other classes of TEP were recently discovered in insects: the insect TEP (iTEP) (Lagueux et al., 2000) and the macroglobulin/complement-related (Mcr) (Stroschein-Stevenson SL, 2006), which constitute a third subfamily inside TEP superfamily, based on phylogenetic analysis (Shokal and Eleftherianos, 2017).

TEP proteins are key components of innate immunity: Complement factors are deposited on the pathogen surface, enhance phagocytosis (opsonisation), can recruit phagocytic cells at sites of infection (chemotaxis), and are capable of direct pathogen lysis; while A_2_Ms are pan-protease inhibitors that sequester pathogen proteases, inhibiting their activity, and promoting their clearance (Williams and Baxter, 2014;Baxter et al., 2017).

TEPs seem to appear early in animal evolution: members of this family are present in a wide variety of invertebrate species: in arthropods (Levashina et al., 2001;Blandin and Levashina, 2004;Buresova et al., 2011;Sekiguchi et al., 2012;Urbanová et al., 2015); crustaceans (Wu et al., 2012;Li et al., 2017); cnidarians (Fujito et al., 2010) and mollusks (Zhang et al., 2007;Yazzie et al., 2015).

In invertebrates, TEPs have primarily been studied in *Anopheles gambiae* and *Drosophila melanogaster* (Vierstraete et al., 2004;Blandin et al., 2008). The *Anopheles gambiae* genome encodes 19 homologues of invertebrate TEP (iTEP) (AgTEP1-19), of which AgTEP1 is the best functionally characterised (Blandin et al., 2008) and structurally the only crystallised TEP from invertebrates (Baxter et al., 2007). AgTEP1 is reported to play an opsonin role in the phagocytosis of Gram-negative and Gram-positive bacteria (Levashina et al., 2001;Yassine and Osta, 2010;Yassine et al., 2012). AgTEP1 also has the capacity to bind to the plasmodium parasite surface to promote its melanisation (Blandin et al., 2008), and thus plays an essential role in decreasing the plasmodium ookinete load in a mosquito’s gut (Smith et al., 2015). The *Drosophila melanogaster* genome encodes 6 homologues of iTEP (DmTEP1-6) (Blandin and Levashina, 2004). Except DmTEP5, DmTEPs are expressed in immune tissues, i.e. hemocytes or fat body, and are up-regulated after immune challenges with bacteria or yeasts (Wertheim et al., 2005;Bou Aoun et al., 2010;Arefin et al., 2014). DmTEP2, DmTEP3 and DmTEP6 bind to Gram-negative, Gram-positive bacteria and fungi respectively, and play an opsonin role to promote the phagocytosis (Stroschein-Stevenson et al., 2006;Bou Aoun et al., 2010). Further, a mutant fly line lacking the four immune-inducible TEPs (TEP1-4) showed lower survival ability following Gram-positive bacteria, fungi or parasitoid wasp immune challenges. This mutant fly line also presented a reduced Toll pathway activation upon microbial infection, leading to a reduced antimicrobial peptide gene expression and thus a less efficient phagocytic response (Dostálová et al., 2017). Interestingly, another close related protein Mcr has been identified for its ability to bind the cell surface of *Candida albicans* to promote its phagocytosis (<u>Shannon L Stroschein-Stevenson</u> et al;2006). This protein, characterised by the lack of the critical cysteine residue within the atypical thioester site (ESGEQN), is not processed during the interaction suggesting that the full-length protein is the active recognition form (Stroschein-Stevenson, 2006 #23). Besides its role in immunity, Mcr is involved in autophagy regulation *via* the Draper immune receptor (Lin, 2017 #66) and in septate junction formation (Batz, 2014 #68; Hall, 2014 #67). More recently, a TEP has been identified in the shrimp *Litopenaeus vannamei* (LvTEP1), and its protective role against both Gram-positive and Gram-negative bacteria and viruses was highlighted by a knockdown approach (Li et al., 2017). Also, the potential immune role of the *Chlamys farreri* TEP (CfTEP) has been suggested as CfTEP transcripts expression is induced following bacterial challenges, while CfTEP protein undergoes an apparent cleavage in a similar manner as the vertebrate’s C3 complement (Xue et al., 2017).

Here we report the potential immune role of TEP protein from the freshwater snail *Biomphalaria glabrata* (BgTEP). In the last decade, *Biomphalaria glabrata* has attracted attention due to its medical and epidemiological importance as a vector for Schistosomiasis disease (King et al., 2005). Authors have invested considerable effort to investigate the molecular interactions and compatibility (susceptibility/resistance status) between *B. glabrata* and its parasite *Schistosoma mansoni* (Mitta et al., 2012;Portet et al., 2017) to help in the discovery of new ways to prevent and/or control Schistosomiasis disease in the field (Tennessen et al., 2015). Co-immunoprecipitation (CoIP) experiments, using the previously characterised *S. mansoni* polymorphic mucins (SmPoMucs) as bait (Roger et al., 2008a;Roger et al., 2008b;Roger et al., 2008c), enabled us to identify putative SmPoMuc-snail interacting immune receptors. An immune complex that associates 3 partners has been characterised: 1) SmPoMucs parasite molecules, 2) Fibrinogen-related proteins (FREP) which is a highly diversified lectin family from snail hemolymph secreted by hemocytes and considered as pathogen recognition receptors (Adema et al., 1997;Zhang et al., 2004;Pila et al., 2017), 3) and a third partner, the newly identified *B.glabrata* Thioester-containing Protein (TEP) named BgTEP (Moné et al., 2010). The BgTEP is characterised by a stretch of cysteine residues at the C-terminal part and a highly conserved thioester motif (GCGEQ) of TEP family (Moné et al., 2010). Interestingly, BgTEP was firstly characterised as an alpha macroglobulin proteinase inhibitor by Bender RC and Baynes CJ, who identified the very first N-teminal amino acids by Edman degradation. They demonstrated that BgTEP is able to inhibit in a methylamine-dependent manner, the activity of cysteine proteinases produced by *S. mansoni* miracidium and sporocysts.

In the present study, we show that structural modelling prediction displays a strong similarity between BgTEP and AgTEP1, while the expression in different tissues reveals a wide distribution, showing a high abundance in snail hemocytes (circulating immune cells). Moreover, BgTEP displays a capacity to interact with diverse intruders, followed by a biochemical processing of the protein. Among hemocyte cells, BgTEP is secreted by a range of blast-like cells which are not phagocytic cells, but which are involved in the formation of the capsule surrounding the parasite. All of these results suggest that BgTEP may play a role in innate immune response against pathogens by assuming an opsonin function as evidenced in arthropods.

## Materials & Methods

### Ethical statements

Our laboratory holds permit # A66040 for experiments on animals, which was obtained from the French Ministry of Agriculture and Fisheries and the French Ministry of National Education, Research, and Technology. The housing, breeding and care of the utilised animals followed the ethical requirements of our country. The experimenter possesses an official certificate for animal experimentation from both of the above-listed French ministries (Decree # 87-848, October 19, 1987). The various protocols used in this study have been approved by the French veterinary agency of the DRAAF Languedoc-Roussillon (Direction Régionale de l’Alimentation, de l’Agriculture et de la Forêt), Montpellier, France (authorisation # 007083).

### Biological materials

The albino Brazilian strain snails (Recife, Brazil) were exposed to several microbes, Gram-negative bacteria culture of *Escherichia coli*, Gram-positive bacteria culture of *Micrococcus luteus* and yeast culture of *Saccharomyces cerevisiae.* Snails were also exposed to one Guadeloupean parasite strain of *Schistosoma mansoni* (le Lamentin, Guadeloupe). This last interaction was chosen for its incompatibility (Galinier, 2017 #70), which means, the snail immune response is efficient and an encapsulation around the parasite is observed.

### Antibody production

We used two types of antibody. The first is a purified polyclonal anti-BgTEP antibody produced in rabbits and named: anti-BgTEP-PEP. The rabbit immune serum was purified on an affinity column raised against immunogenic peptide produced by Eurogentec. The sequence of the peptide is H2N – SSY GSK SFR PDT NIT C – CONH2. The second antibody is a purified polyclonal anti-BgTEP antibody produced in chickens and named: anti-BgTEP-RP. A truncated recombinant protein which corresponds to the C-terminal part of BgTEP from nucleotides 1,036 to 1,445 corresponding to the amino-acid sequence [_1036_LIDRQ-CLNCCP_1445_], was produced in *E. coli* and purified on nickel column (Agrobio manufactures).

### Genomic, Proteic and Structural Organisation of BgTEP

The cDNA sequence of BgTEP, was BLASTed against VectorBase to characterise the exon-intron structure (data not shown). The 5’UTR and 3’UTR ends were determined using data of the Brazilian *B. glabrata* transcriptome (Dheilly et al., 2015) Prediction of the BgTEP threedimensional (3D) structure and alignment with AgTEP1 (2PN5 published in 2015 (Baxter et al., 2007) were performed using the I-Tasser and TM-align servers (Zhang and Skolnick, 2005;Zhang, 2008). The 3D structure was obtained by multiple threading using the I-Tasser server (available online), which combines two protein structure prediction methods: threading and *ab initio* prediction (Roy et al., 2010). Structural similarities between the functional domain of AgTEP1 and BgTEP were determined by calculating a TM-score. A TM-score greater than 0.5 reveals significant alignment, whereas a TM-score less than 0.17 indicates a random similarity.

### Phylogenetic Organisation of TEP family members

Homologous sequences were identified using BLAST searches against the GenBank non redundant database (Bethesda, USA). For phylogenetic analyses, multiple protein sequence alignments were performed with ClustalW using the BLOSUM62 substitution matrix model. The neighbor-joining method (Poisson substitution model; uniform substitution rate; gaps/missing data treatment: pairwise deletion) was used to generate the phylogenetic tree, in order to cluster the different protein families, and to determine BgTEP position. Neighbor-joining method was chosen as full length protein sequences of varying size and harbouring different conserved domains were used to construct the tree. A bootstrap analysis of 2000 replications was carried out to assess the robustness of the tree branches. A total of 125 full length sequences from TEP superfamily proteins (data not shown) were chosen to construct the tree based on the best BLAST matches against the NCBI database. The phylogenetic analysis was performed using MEGA 5 software (Tamura et al., 2011). Besides, the search for a C-terminal transmembrane domain was performed to distinct CD109 from iTEP proteins, using the online transmembrane topology and signal peptide predictor Phobius from the Stockholm bioinformatic center (http://phobius.sbc.su.se/). The result of this search was used to classify invertebrate CD109-like proteins and iTEP in different subgroups in line with the phylogenetic tree obtained.

### BgTEP interactome and immunoblotting approaches

<u>*B. glabrata plasma preparation:*</u> The hemolymph was extracted from the head-foot according to previously described procedures (Sminia and Barendsen, 1980). After recovering the hemolymph, a first centrifugation for 5 min at 5,000 g at 4°C was performed to eliminate hemocytes then, an ultracentrifugation for 2.5h at 40,000 g at 4°C leads to hemoglobin elimination. The ultracentrifugated plasma was only used for the study of the BgTEP profile in naive snail and to test the specificity of the different antibodies

<u>*Interactome samples preparation:*</u> 2 ml of fresh hemolymph were recovered from naive snails and centrifuged for 5 min at 5,000 g to eliminate hemocytes. In parallel, live microbe samples were recovered, centrifuged at 5,000 g for 5 min and washed with CBSS (NaCl 2.8 g/L, KCL 0.15 g/L, Na_2_HPO_4_ 0.07 g/L, MgSO4.7H_2_O 0.45 g/L, CaCl_2_.2H_2_O 0.53 g/L, NaHCO_3_ 0. 05 g/L) and kept at the bottom of the tube prior to use. Depending on the intruder, samples were recovered from 500 μL of bacteria-culture of *Escherichia coli* and *Micrococcus luteus* (1.2×10^7^ part/mL), 500 μL of yeast-culture of *Saccharomyces cerevisiae* (7.5×10^5^ part/mL) and 500 miracidia (snail infective stage of *S. mansoni*) or 500 primary sporocysts (first snail-stage of *S. mansoni*). Two interactome conditions were then tested for each adopted intruder: intruders exposed either to (i) the host cell-free hemolymph fraction or to (ii) CBSS that mimics the internal host environment. Samples were incubated at 26°C (temperature of snail environment) for 30 min, to observe a rapid interaction between the BgTEP and the intruders, or 3 hours to observe potential processing of BgTEP. Intruder samples incubated with hemolymph or CBSS buffer were then washed with CBSS as detailed above and pellet intruders were extracted using Laemmli buffer (Bio-Rad) with *β*-mercaptoethanol. The biological material was used to study the BgTEP profile after interaction between the cell-free hemolymph and intruders.

<u>*Western-blot:*</u> 8 μL of snail ultracentrifugated plasma or 10 μL of interactome samples were run on a 7.5% SDS-polyacrylamide precast gel (Mini Protean TGX Precast Gel Bio-Rad) for bacteria and yeast samples and 12% SDS-polyacrylamide precast gel (Mini Protean TGX Precast Gel Bio-Rad) for parasite samples and transferred onto a 0.2 μm PVDF membrane with Trans-Blot Turbo Transfer Pack (Bio-Rad). After saturation during 1h at 37°C in TBSTM (1x TBS (500 mM Tris-HCl, 1.5 M NaCl, pH 7.5), 0.05% Tween-20, 5% non-fat milk), the protein blots were incubated for 1.5h at room temperature in TBSTM, with a 1:200 dilution of anti-BgTEP-PEP antibody for the western-blot on ultracentrifuged naive snail plasma and a 1:2000 dilution of anti-BgTEP-RP antibody for the interactome experiments. The blots were washed 3 times with TBST, and further incubated with 1:4000 dilution of the manufactured horseradish peroxidase-conjugated goat anti-rabbit IgG antibody (Pierce) for the western-blot on naive snail plasma, and 1:4000 dilution of the manufactured horseradish peroxidase-conjugated goat antichicken IgY antibody (Southern Bioteck) for the interactome experiments, in TBSTM for 70 min at room temperature. Blots were finally washed 3 times with TBST, and then revealed in the presence of an enhanced chemiluminescent subtract (Super Signal West Pico Chemiluminescent Substrate, Thermo Scientist). These experiments were performed at least three times independently and one representative blot is shown.

### BgTEP immunolocalisation in hemocyte populations

The hemolymph was extracted from the head-foot according to previously described procedures (Sminia and Barendsen, 1980). The hemocytes were plated for 1h on polystyrene chamber-slides. Hemocytes were fixed using 4% paraformaldehyde for 10 min, followed by a cell permeabilisation step with 0.01% Triton X-100 and 3% BSA for 20 min. Cells were incubated with a 1:100 dilution of anti-BgTEP-PEP for 1h followed by a 1:1000 dilution of the manufactured fluorescent goat anti-rabbit IgG antibody (Thermo Fisher Scientific – Alexa Fluor 594) for 50 min. After the BgTEP labelling, the actin was labelled with (Alexa Fluor 488) Phalloidin (Thermo Fisher Scientific) for 20 min and the cell nucleus with Dapi (Biotum) for 30 sec. Observations were performed by confocal microscopy using a Zeiss LSM 700 microscope at 405, 488 or 555 nm). These experiments were performed at least ten times independently with the anti-BgTEP-PEP antibody. Negative controls were done to confirm the specificity of the anti-BgTEP-PEP antibody *i.e.* using either secondary antibody alone or preincubating the anti-BgTEP-PEP antibody with the immunogenic peptide. No cell labelling was observed.

### Phagocytosis assay

5 μL of *Saccharomyces cerevisiae* conjugated with an Alexa Fluor 488 (Invitrogen, BioParticles Z23373) *i.e.* 1.10^6^ were injected into snails. The hemolymph was recovered 3h post-injection. The hemocytes were plated on chamber-slides and prepared as described above. Two types of labelling were performed. Either, an alexa fluor 594 phalloidin was used to label actin of all immune cells or the anti-BgTEP-PEP antibody detected using Alexa Fluor^®^ 594 conjugated goat anti-Rabbit IgG secondary antibody to reveal BgTEP positive cells. The internalised green yeasts in different hemocytes were monitored by several focal observations using the Zeiss LSM 700 confocal microscope. These experiments were performed at least three times independently.

### Fluorescent staining and flow cytometry method

The hemolymph was extracted as described above. Three biological replicates (pools of 15 snails per replicate) were performed. Hemocytes were fixed in suspension using 4% paraformaldehyde for 10 min, followed by a cell permeabilisation step with 0.01% Triton X-100 and 3% BSA for 20 min. Cells were incubated with a 1:100 dilution of anti-BgTEP-PEP for 1h followed by a 1:1000 dilution of the manufactured fluorescent goat anti-rabbit IgG antibody (Thermo Fisher Scientific – Alexa Fluor 594) for 50 minutes. Hemocyte population was profiled along using Side Scatter Channel (SSC) and Forward Scatter Channel (FSC) to estimate cell granularity and cell size, respectively. The PE channel was used to detect light emitted from Alexa Fluor 594 dye conjugated to the secondary antibody. The flow cytometry was performed using a FACSCanto from BD Biosciences (RIO Imaging Platform, Montpellier, France). The threshold was determined according to the PE channel and the FSC parameter. Note that the largest cells tend to slightly autofluoresce. For each sample, about 10,000 events were counted. The results were analysed with the FlowJo V 10.0.8 software.

### BgTEP expression analysis

<u>*RNA extraction and Quantitative RT-PCR protocol:*</u> Snail total RNA was extracted with TRIzol reagent (Sigma Life Science) according to manufacturer’s instructions and subsequently reverse transcribed to first strand cDNA using Maxima H Minus First Strand cDNA Synthesis Kit with dsDNase (Thermo Scientific) according to manufacturer’s instructions.

Real-time RT-PCR analyses were performed with the LightCycler 480 System (Roche) in a 10 μL volume comprising 2 μL of cDNA diluted to 1:200 with ultrapure-water, 5 μL of No Rox SYBR Master Mix blue dTTP (Takyon), 1 μL of ultrapure-water and 10 μM of each primer. The primers used for the RT-QPCR are TEP-R: ACCATTAGATCCACTGGAAGATA TEP-F: CTGACTTACCCTCGCTC for BgTEP, and S19-R: CCTGTATTTGCATCCTGTT S19-F: TTCTGTTGCTCGCCAC for S19 ribosomal protein gene used as housekeeping gene. The two primer couples have been tested to determine the exponential and efficiency of the PCR reaction. The cycling program is as follows: denaturation step at 95°C for 2 min, 40 cycles of amplification (denaturation at 95°C for 10 sec, annealing at 58°C for 20 sec and elongation at 72°C for 30 sec, with a final elongation step at 72°C for 5 min. QPCR was ended by a melting curve step from 65°C to 97°C with a heating rate of 0.11°C/sec and continuous fluorescence measurement. For each reaction, the cycle threshold (Ct) was determined using the 2^nd^ derivative method of the LightCycler 480 Software release 1.5 (Roche). PCR experiments were performed in triplicate (technical replicates) from four biological replicates. The mean value of Ct was calculated. Corrected melting curves were checked using the Tm-calling method of the LightCycler 480 Software release 1.5.

<u>*Tissue recovery:*</u> Tissues were collected from 9-10 snails under binocular microscope dissection. Albumen gland, head-foot, hepato-pancreas, and ovotestis were recovered. For hemocytes recovery, the hemolymph of 50 snails was collected, and cells were recovered after centrifugation at 10,000g for 10 min. The relative expression of BgTEP was calculated with the E-method which accounts the efficiency of both couple of primers. Results were normalised with respect to a housekeeping gene the S19 ribosomal protein gene, as previously described (Galinier et al., 2013). Statistical analysis was done by a one way ANOVA test followed by a post-hoc Games-Howell pairwise comparison test.

<u>*Infection by multiple intruders of whole snails:*</u> Contact with Gram-positive and Gram-negative bacteria and yeast were established according to previously described procedures (Deleury et al., 2012). Briefly snails were bathed with 10^8^ part/mL of microbes for 1h, then snails were washed. For the parasite infection, each snail was exposed for 6h to 10 miracidia in 5 mL of pond water. For each intruder stimulation, 36 snails were used: four independent replicates using a pool of 3 snails were performed at 3 time points (6h, 12h and 24h after stimulation). As control condition, 4 replicates using a pool of 3 snails were recovered for the evaluation of the basal expression of BgTEP. The relative expression of BgTEP were normalised with respect to a housekeeping gene the S19 ribosomal protein gene, and BgTEP expression was compared to non-exposed snails. The significant difference in BgTEP expression was evaluated based on ΔCt values, according to a Kruskal-Wallis test and followed by a Dunn Post-hoc test.

PCR experiments were performed in triplicate (technical replicates) from four biological replicates. The mean value of Ct was calculated. Corrected melting curves were checked using the Tm-calling method of the LightCycler 480 Software release 1.5.

### *In situ* histological localisation of BgTEP

<u>*Snail infection:*</u> Snail were infected with the parasite strain as previously described, briefly each snail was exposed for 6h to 10 miracidia in 5 mL of pond water.

<u>*Immunocytochemistry procedures:*</u> Snails were fixed in Halmi’s fixative (4.5% mercuric chloride, 0.5% sodium chloride, 2% trichloroacetic acid, 20% formol, 4% acetic acid and 10% picric acid-saturated aqueous solution). Embedding in paraffin and transverse histological sections (10-μm) were performed. The slides were stained using Anti-BgTEP-PEP antibody according to a previously developed protocol (Roger et al., 2008a).

Slides re-hydration was performed in serial toluene, 95%, 70%, and 30% ethanol and finally PBS bathing (Saline Phosphate Buffer: pH 7,4-7,5; 8,41 mM Na_2_HPO4; 1,65 mM Na_2_H2PO_4_; 45,34 mM NaCl; H20 milliQ q.s.p.). The slides were immersed in a permeabilising PBS solution containing 0.5% Triton X-100 during 15 min. A saturation step was performed in PBS buffer containing 1% gelatine hydrolysate (Bellon, France), 1% normal goat serum (NGS, Sigma) and 0.1% NaN3 (Sigma) during 1h at room temperature. Slides were then successively incubated with the Anti-BgTEP-PEP antibody dilution 1:100 for 1h30 at room temperature and with an Alexa Fluor 594 anti-rabbit IgG (Thermo Fisher Scientific) diluted 1:1000 for 1h at room temperature. Slides were mounted in Vectashield and stored in dark at 4°C. The slide observation was carried out by epifluorescence and light microscopy using a Zeiss axioscope 2 microscope (Carl Zeiss AG) AG) and a Leica DC350FX camera (Leica).

## Results

### Organisation of BgTEP

BgTEP exhibits all the characteristics of known iTEP family members, including a signal peptide for secretion, several predicted N-glycosylation sites, the canonical thioester motif (GCGEQ), the complement component domain (pfam PF07678), the Alpha2 Macroglobulin receptor binding domain (pfam PF07677) and numerous cysteins such as the 6 C-terminal ones which are a signature of iTEPs (Moné et al., 2010). At the genomic level, BgTEP is composed of 37 exons spreading on 10 scaffolds of the *Biomphalaria glabrata* genome assembly (Adema et al., 2017) (Figure 1A). Exon-intron boundaries were conserved along the sequence (data not shown). At the secondary structure level, BgTEP protein possesses an N-terminal alpha helix characteristic of signal peptide for secreted proteins, like other iTEPs. As expected, BgTEP is composed of 8 MG (Macroglobulin) domains, a succession of several *β*-sheets related to the fibronectin type III (fnIII) domains, with insertions of a LNK domain nested into MG6 and a CUB domain (*β*-sheet domain), and of a TED domain (*α*-helical thioester domain) between MG7 and MG8 (Figure 1 A).

**Figure 1:**
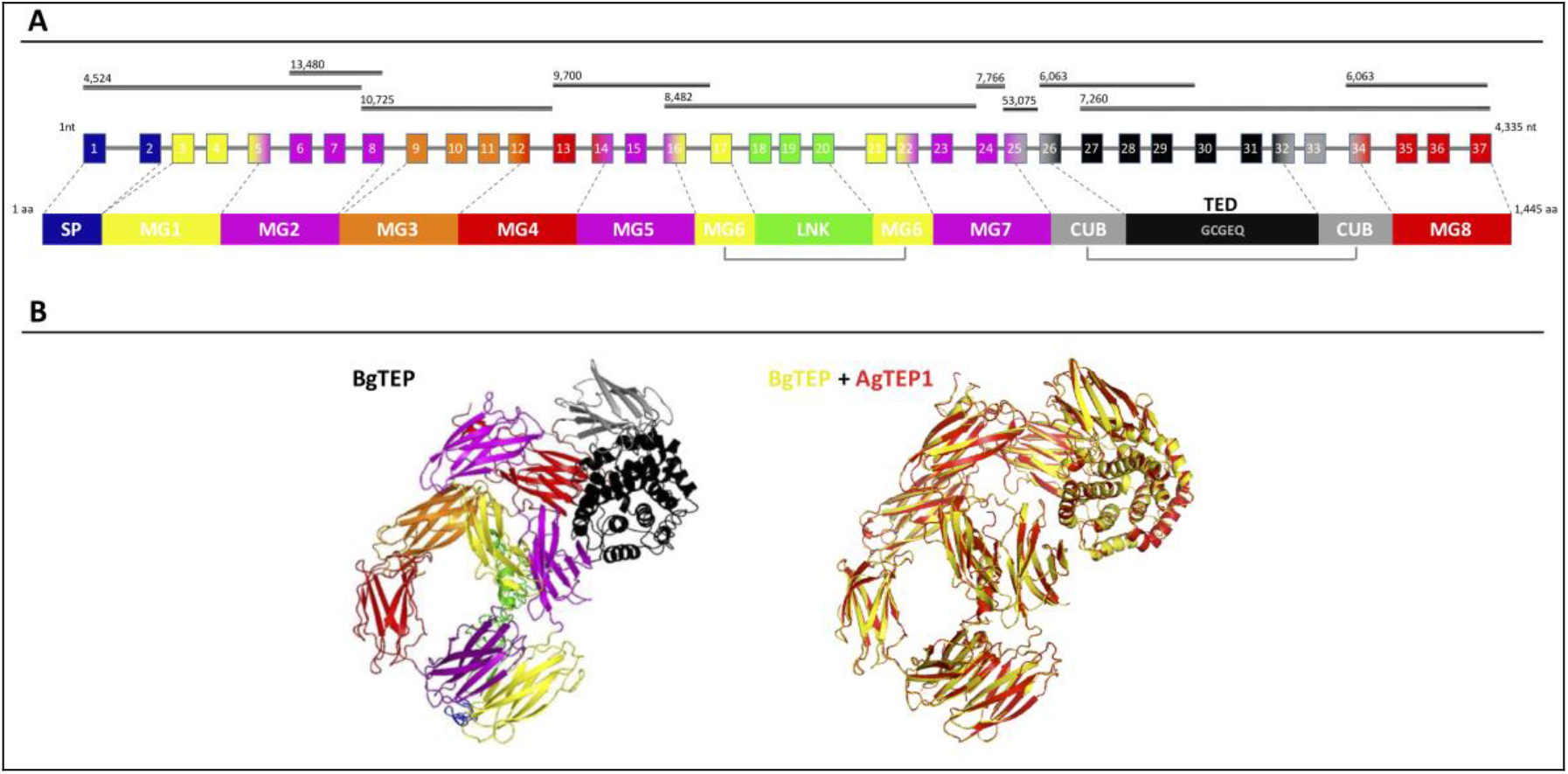
BgTEP organization. A. Schematic representation of BgTEP sequence organisation. The top black lines correspond to Biomphalaria glabrata genome scaffolds assigned with their corresponding scaffold numbers. The centre part is a schematic of the exon/intron organisation of BgTEP gene. The squares represent exons, the lines for introns, the number corresponds to the exon number, and finally their associated colour corresponds to the different protein domain encoded that mapped on 3D structure. The lower part corresponds to the schematic domain arrangement of BgTEP protein. The signal peptide (SP) domain is represented in blue followed by macroglobulin domain 1 (MG1) in yellow, MG2 in magenta, MG3 in orange, MG5 (magenta), MG6 (yellow), linker domain (LNK) (green), MG7 (magenta), CUB (grey), thioester domain (TED) with conserved motif (GCGEQ) in black andMG8 in red. B. Overview of three-dimensional BgTEP structure predicted by I-Tasser software, using Anopheles gambiae TEP1 (AgTEP1) as reference. The coloured domains of the BgTEP protein match the color of the corresponding encoding exon. Despite a low similarity at the amino-acid level (29%), a high conserved spatial conformation (TM Score = 0.98) was observed between BgTEP (yellow) and TEP1 of Anopheles gambiae (red).

The BgTEP protein’s tertiary structure was also investigated with the structure prediction software I-Tasser (Figure 1B), using AgTEP1 protein from *Anopheles gambiae* as reference structure. The score obtained for this prediction is highly significant (TM Score = 0.98), indicating the robustness of the prediction, as well as a structural alignment conservation with the AgTEP1 reference despite a low similarity at the amino-acid level (29%) (Figure 1B).

### Phylogenetic analysis of BgTEP

Since the first identification of BgTEP in 2010 (Moné et al., 2010), many new members of the TEP superfamily have been discovered, including several from the phylum *Mollusca*. In response to these recent discoveries, we conducted a new phylogenetic analysis to position BgTEP among the main TEP family members. To do this we used 125 amino acid sequences of full length TEP superfamily proteins (data not shown), including complement-like factors, macroglobulin complement-related proteins (MCR), A_2_M, complement 3 and pregnancy zone protein-like A_2_M domain-containing 8 (CPAMD8), CD 109 glycoproteins and iTEP proteins, to construct a phylogenetic tree with the neighbour-joining method (Figure 2). Phylogenetic analysis confirms the localisation of BgTEP within the group of iTEPs. This analysis segregates the proteins into three major groups; the complement component group which includes complement and complement-like factors as well as MCR (Figure 2, blue shades), the A_2_M group comprising A_2_M and CPAMD8 (Figure 2, red shades), and the group formed by cell surface TEP (CD 109) and iTEP (Figure 2, green shades). However, the analysis reveals several subgroups within the iTEP and CD 109 group. Vertebrate and invertebrate CD 109 proteins form two distant subgroups, while invertebrate TEP and invertebrate CD109 form two closely related subgroups. CD109-like proteins from invertebrate may be distinguished from iTEP proteins by the presence of a putative C-terminal transmembrane domain (Wu et al., 2012), as reported for the vertebrate GPI-anchored CD 109 proteins (Solomon et al., 2004). Moreover, BgTEP is included in an additional well-supported subgroup constituted by both CD 109 and iTEP from mollusk species, reflecting a degree of similarity in the sequences of TEP proteins from the phylum Mollusca. The mix of CD 109 and iTEP in the same subgroup is likely the result of automatic – and not manual – corrected annotation from Genbank without further characterisation.

**Figure 2:**
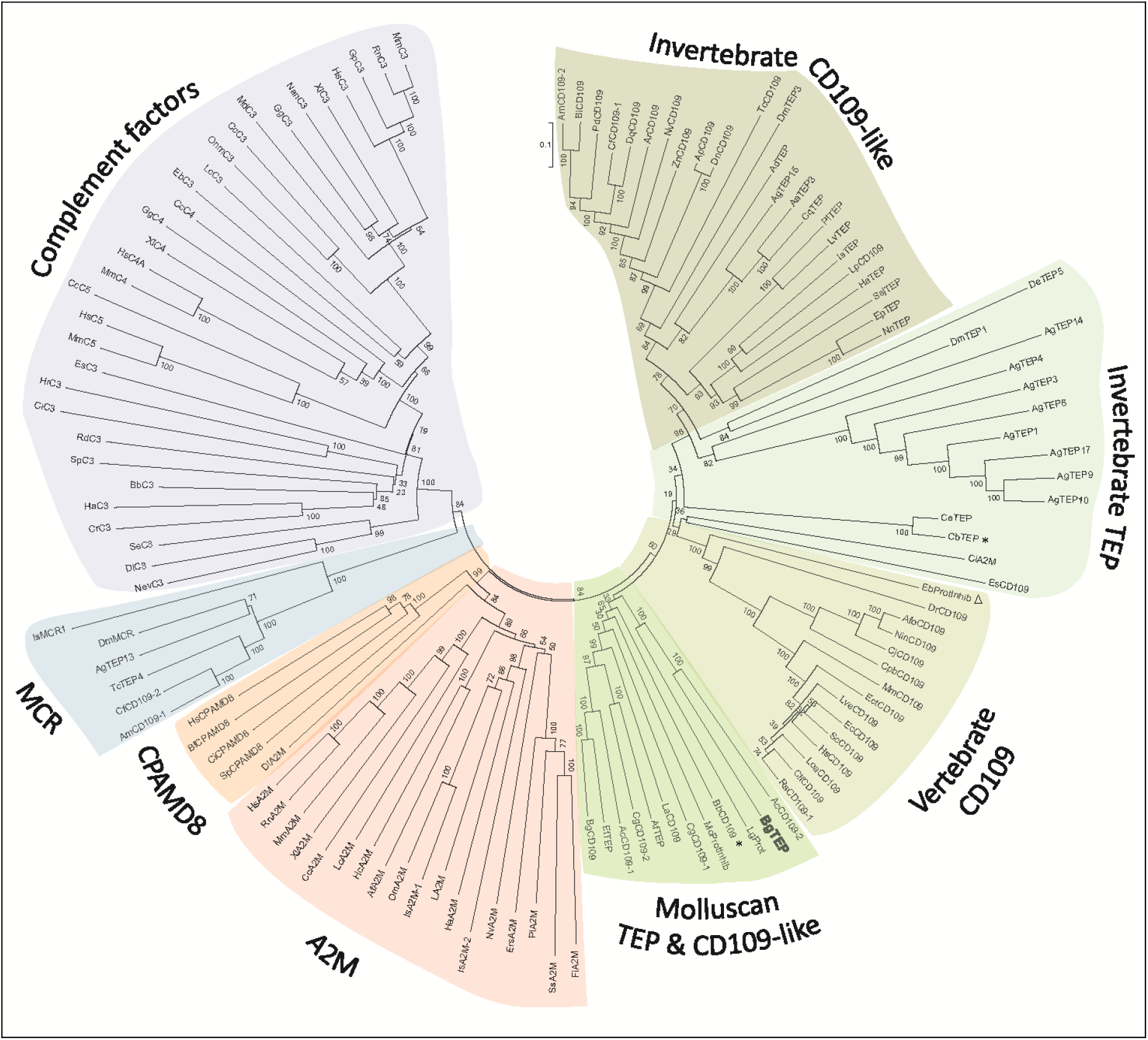
Phylogenetic Tree of TEP superfamily. Phylogenetic analysis of the TEP superfamily full length protein sequences from 125 members. Complement factor groups are coloured in blue shades, alpha-2 macroglobulin groups in red shades, and iTEP and CD 109 groups in green shades. BgTEP is indicated in bold type. A bootstrap analysis of2000 replications was carried out on the tree inferred from the neighbour joining method and the values are shown at each branch of the tree.

### BgTEP Expression in snail tissues

A dissection of different organs (hemocytes, ovotestis, head-foot, hepatopancreas and albumen gland) of *B. glabrata* was conducted to analyse tissue distribution of BgTEP by quantitative RT-PCR (Figure 3). BgTEP was expressed in all tissues examined with different levels; a high expression is observed in hemocytes, ovotestis and head-foot, whereas a lower expression is observed in hepatopancreas and albumen gland (Figure 3).

**Figure 3:**
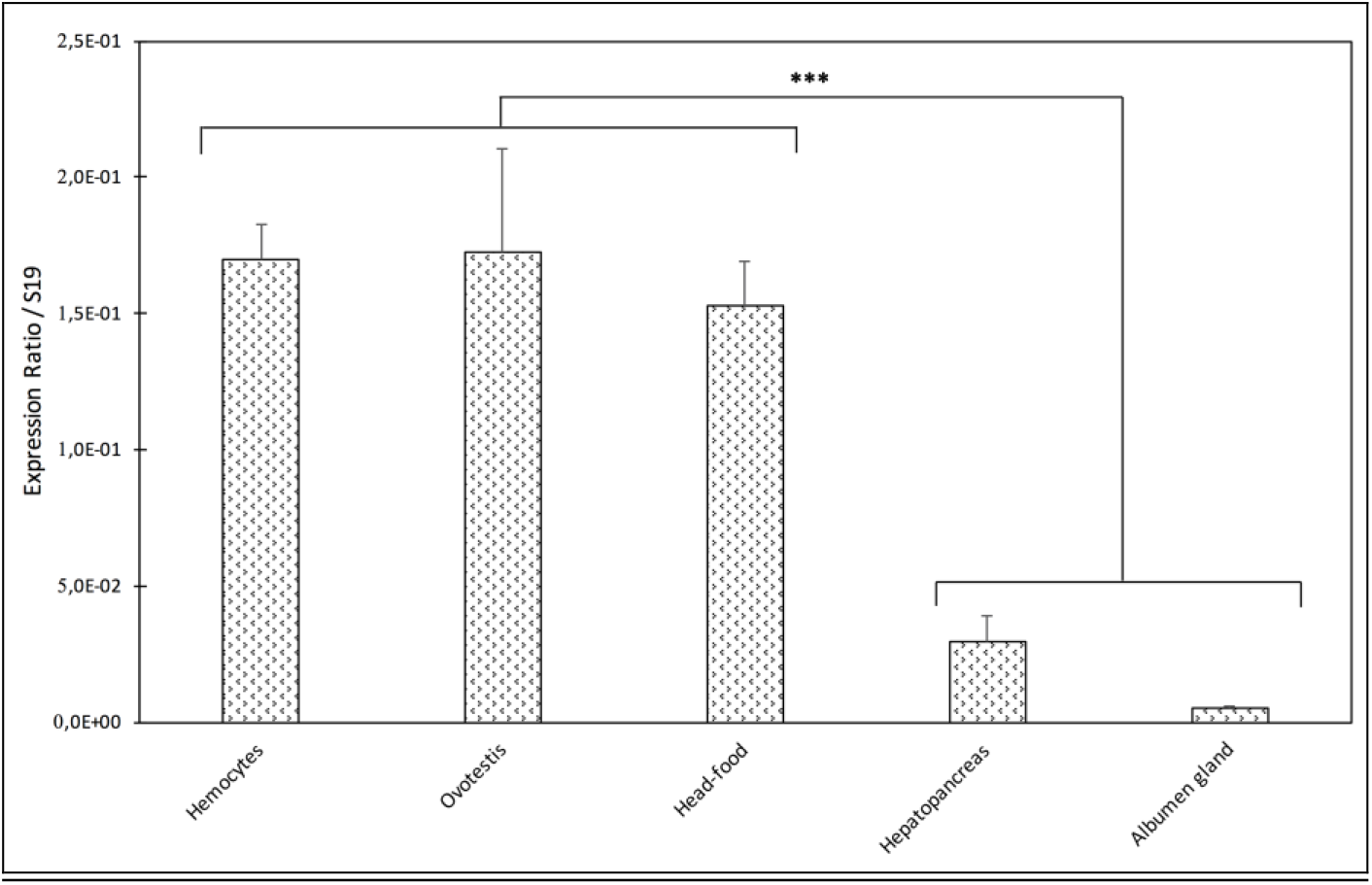
Tissue Expression of BgTEP. Transcription profile of BgTEP in different tissues extracted from 9-10 snails. Five snail tissues are dissected: Albumen gland, Hepatopancreas, Head-foot, Ovotestis. Hemocytes were collected from 50 snails. Quantitative RT-PCR was performed on total RNA. Ct-values of the BgTEP transcript were normalised to the transcript level of the reference gene S19. Error bars represent the standard deviation of the ΔCT mean values obtained for each tissue. Asterisks indicate a significant expression difference between mentioned tissues.

A western blot experiment was performed on ultracentrifuged cell-free hemolymph from naïve snails with a polyclonal anti-BgTEP antibody, designed from a C-terminal peptide and called anti-BgTEP-PEP. We detected 3 bands of BgTEP in naive plasma (Figure 4). The first band abundantly detected at approximately 200 kDa corresponds to the full-length BgTEP (Figure 4). Two lower bands (60 and about 30 kDa) were interpreted as BgTEP processed forms, resulting from a fine regulation of proteolysis by endogenous proteases observed also in mosquito plasma (Figure 4).

**Figure 4:**
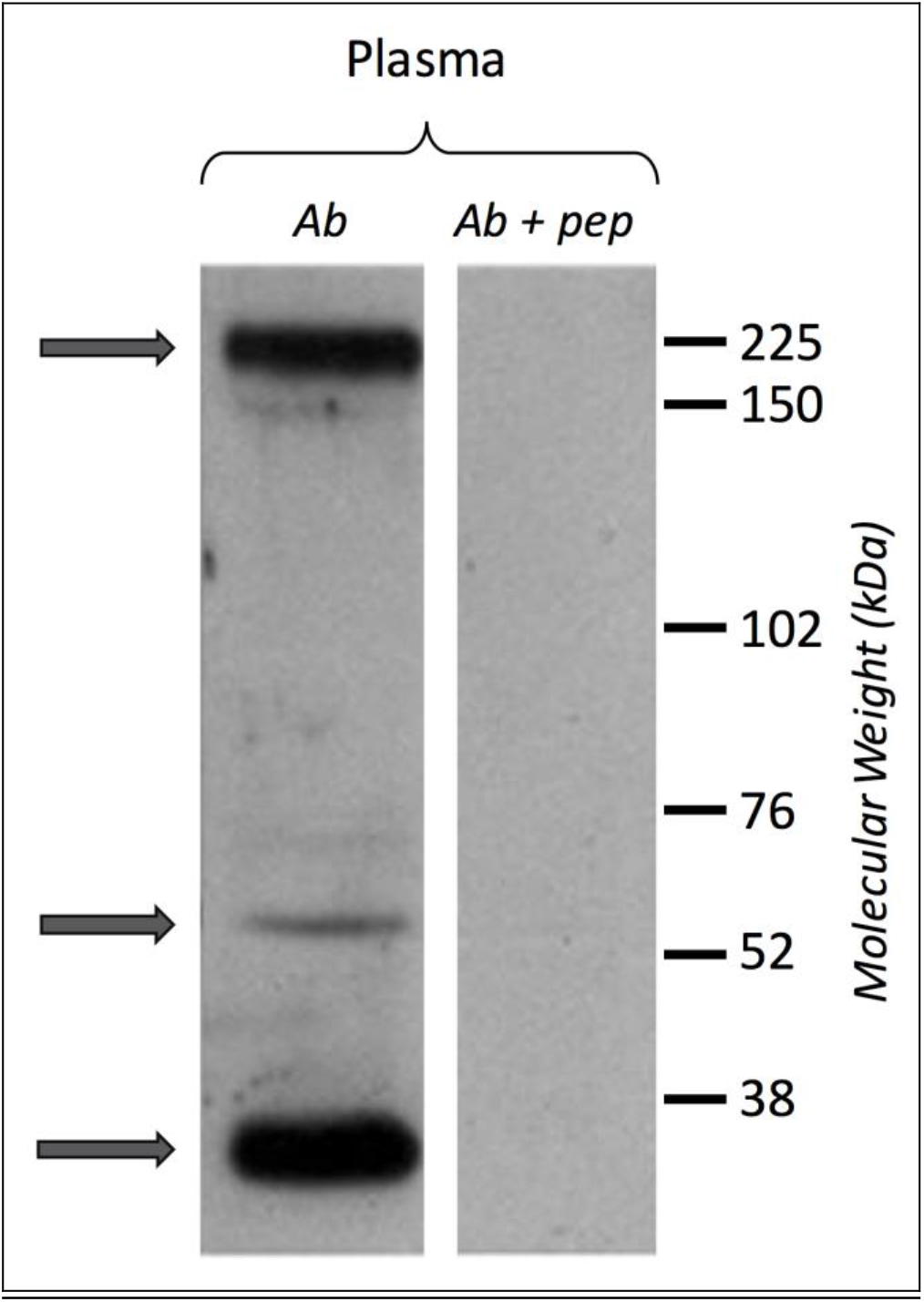
BgTEP Secretion. Presence of BgTEP protein on hemolymphatic cell-free compartment was tested by western blot using an affinity-purified polyclonal antibody (anti-BgTEP-PEP). Ultracentrifuged hemolymph was used to remove hemoglobin. As negative control, an Ab+pep (anti-BgTEP antibody and BgTEP peptide) pre-incubation was performed before incubation with the membrane. Arrows correspond to the full-length and two processed forms of BgTEP protein present in hemolymph from naïve snails.

### BgTEP expression after immune challenges

In some invertebrate models, the iTEP has been suggested to play a role during immune infections, mostly assuming an opsonin function. To investigate this potential function in our *B. glabrata* model, we measured the relative expression ratio by RT-QPCR of BgTEP transcripts in response to different immune challenges (Figure 5). The expression varies greatly depending on intruder immune challenges but also during infection kinetics. Indeed, BgTEP expression is decreased after both *E. coli* and *S. cerevisiae* challenges, while it is up regulated after *M. luteus* and *S. mansoni* stimulations (Figure 5). When challenging with *E. coli*, the expression is rapidly lowered nearly 5 fold, and returns to basal expression level after 24 hours. However, while following the *S. cerevisiae* challenge, BgTEP expression is only slightly decreased two fold from 12h to 24h post-challenge. For the other challenges, we observed a two-fold increase in expression from 6 to 12 hours after *M. luteus* stimulation, while for the *S. mansoni* challenge, the BgTEP expression increases regularly from 6 to 24 hours from 2 to 3.5-fold.

**Figure 5:**
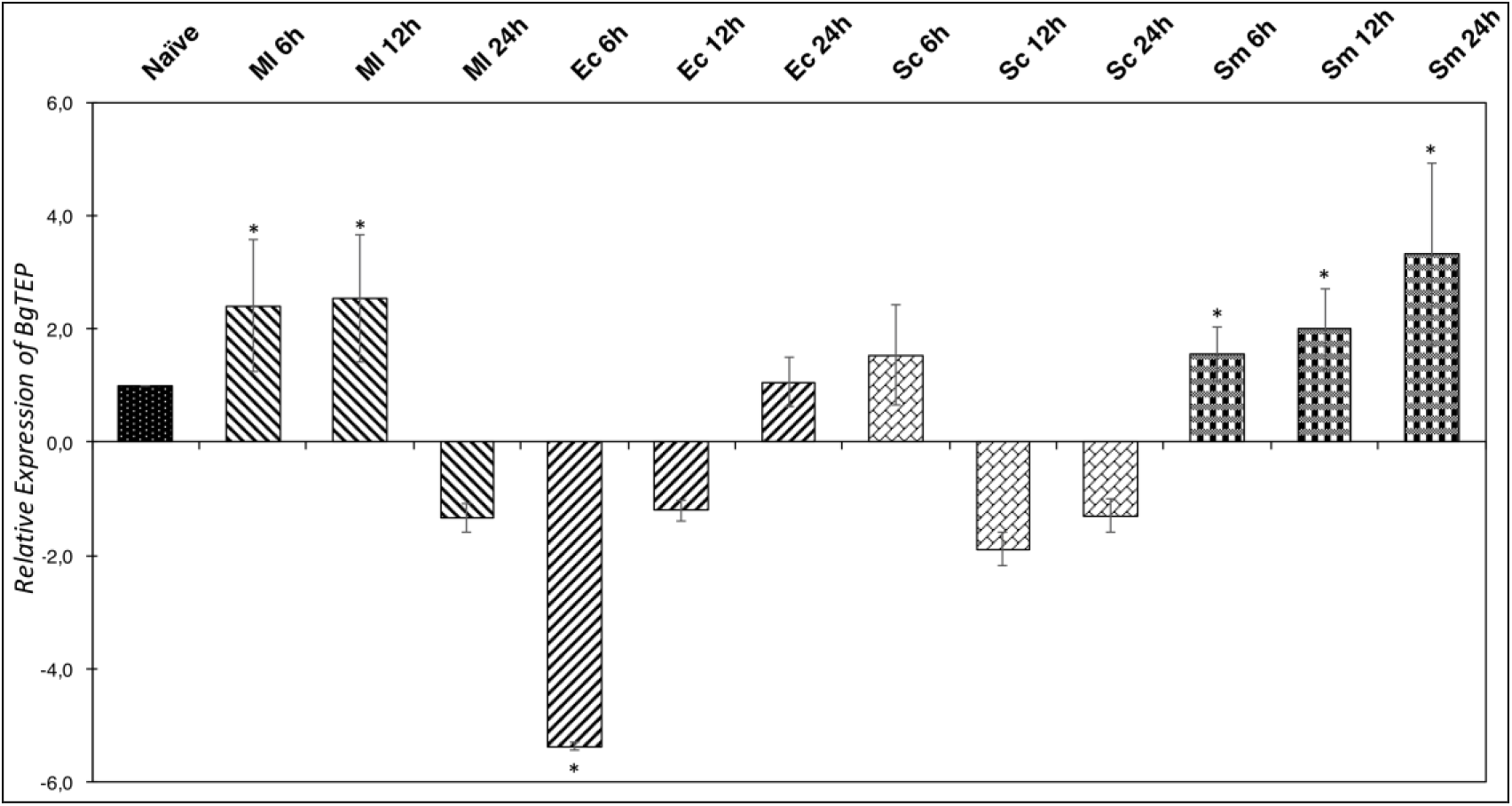
Immune-Inducible Transcription of BgTEP. Quantitative RT-PCR was performed from whole snails exposed to different immune challenges or unexposed. BgTEP expression was monitored at 3 time points (6, 12 and 24 hours) post challenge with M. luteus (decreasing hatching), E. coli (increasing hatching), S. cerevisiae (rectangle) and S. mansoni (square). BgTEP expression was normalised by S19 housekeeping gene expression for each experimental point and compare to the relative expression obtained in non-exposed snails. Error bars represent the standard deviation of the relative quantification values obtained for each kinetic’point. The significant difference in BgTEP expression was evaluated according to a Kruskal-Wallis test followed by a Dunn Post-hoc test. The asterisks indicate a significant difference between non-exposed and exposed snails.

As the number of BgTEP transcripts is regulated following challenges with different intruder types, we wonder if this protein might be involved in pathogen recognition, like in other invertebrate models such as Drosophila or Anopheles, that play an opsonin role in immune recognition response.

### BgTEP capacities to link intruder surface

To investigate the ability of BgTEP to bind to the intruder surface during infections, we performed interactions at 26°C between *B. glabrata* cell-free hemolymph and *M. luteus, E. coli, S. cerevisiae* or *S. mansoni* parasites (Figure 6). As a comparison, intruders alone were incubated at the same time in CBSS buffer (negative control) (Figure 6). The experiments were performed with whole live intruders allowing for two interaction times (30 min and 3h) in order to detect the binding and the potential processing of BgTEP protein during binding kinetics. Once the interactions were complete, intruders were recovered by centrifugation, washed, and bound BgTEP was revealed by western blot using the anti-BgTEP-RP antibody (Figure 6). This antibody raised against the C-terminal part of the BgTEP revealed several bands in positive control, three of which (200, 100 and 50kDa) were more intense (Figure 6A, lane 1).

**Figure 6:**
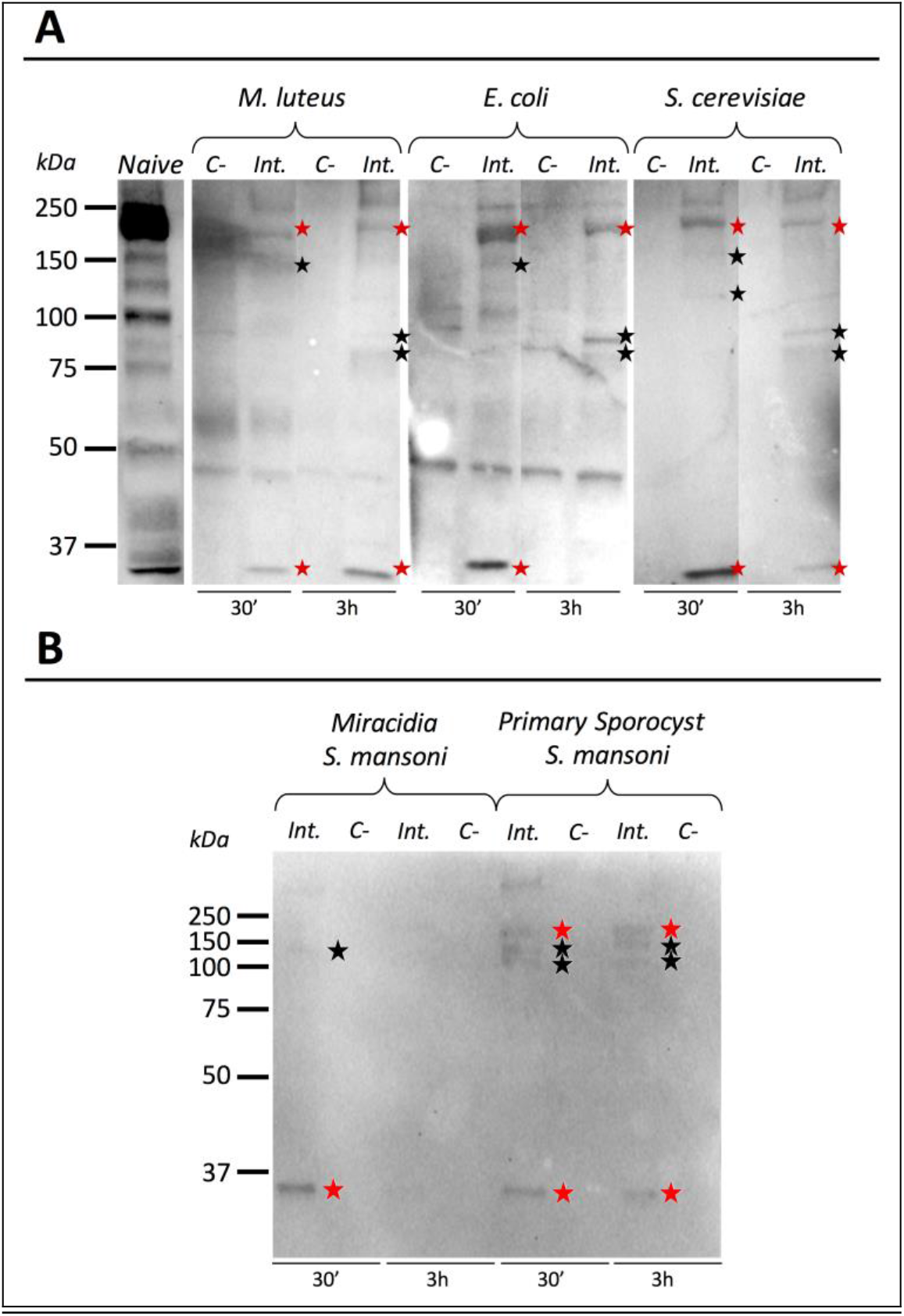
Immunoblotting analysis of BgTEP. Live intruders interaction with B.glabrata cell-free hemolymph was monitored by Western blot using anti-BgTEP-RP antibody. Ultracentrifuged cell-free hemolymph from naïve snail was used as positive control for full-length and processed forms detection of BgTEP. For each interaction condition, negative control (C-) corresponds to the intruder alone, interactome (Int.) corresponds to the interaction between the intruder and B. glabrata cell-free hemolymph. Red stars correspond to putative full-length (200 kDa) and cleaved protein (about 30 kDa) observed in all interactions and black stars indicate bands between 75 and 150 kDa corresponding to BgTEP alternative processed forms. A. Comparison of BgTEP binding and time-dependent processing after 30 min and 3 h of interaction between cell-free hemolymph and M. luteus, E. coli, S. cerevisiae. B. Comparison of BgTEP binding between two developmental stages of S. mansoni (miracidia and primary sporocyst).

Interestingly we demonstrate that the full length of BgTEP (band of 200 kDa) binds all bacteria, yeast (Figure 6A) and metazoan parasites (Figure 6B). Surprisingly, we also observed that a short processed form of TEP (about 30 kDa) present in naïve hemolymph can also bind to all intruders. Other minor bands were detected by the anti-BgTEP-RP antibody from 75 to 150 kDa with *E. coli, M. luteus* and *S. cerevisiae* that differed between 30 minutes and 3 hour interactions (Figure 6A).

We investigated the interaction between BgTEP and two different development-stages of *S. mansoni* parasite; the miracidia, a free-living and swimming larva that infests the host snail, and the primary sporocyst which is the first intra-molluscal stage of the parasite after infestation (Figure 6B). For the interaction with *S. mansoni*, we demonstrate that the full length of BgTEP binds the parasite at the two development-stages, but to a lower extent with miracidia stage (Figure 6B). We also observed that the short processed form of BgTEP (30 kDa) binds the parasite but is no longer detected with miracidia stage at the 3 hour interaction (Figure 6B). Moreover, minor bands were detected from 100 to 150 kDa for the primary sporocyst stage all along the interaction, whereas only a weak band is observed for the miracidia stage (Figure 6B). This discrepancy between the two profiles suggests different BgTEP binding and processing abilities for the two parasite stages.

To conclude, we evidence the binding of BgTEP to all of the tested intruders but also a differential BgTEP intruder-dependent processing that occurs during interaction. The disappearance or the presence of new forms detected on the intruder surface highly suggest that BgTEP likely has a complex modification of its structure with a possible a fine proteolytic regulation required for opsonisation or/and encapsulation.

### BgTEP expression in hemocyte sub-populations

Since BgTEP can bind different intruders and display an opsonin role in other models, we focused on immune activities at the cellular level. First, we performed an immune-labelling experiment upon plated hemocytes to investigate which sub-type expresses BgTEP (Figure 7A). Immunolocalisation shows that not all hemocytes produce and secrete BgTEP and its expression is only restricted to a subset of blast-like cells (Figure 7A). Moreover, to estimate the BgTEP-positive blast-like cell proportion in hemocytes, we performed a cell quantification by flow cytometry (Figure 7B). The BgTEP-positive-cells correspond to about 20% of total hemocytes (Figure 7B). This result has been confirmed by microscopy cell counting. About 50 % of blast-like cells, which represent approximately half of the entire population of hemocytes (data not shown), are BgTEP-positive-cells.

**Figure 7:**
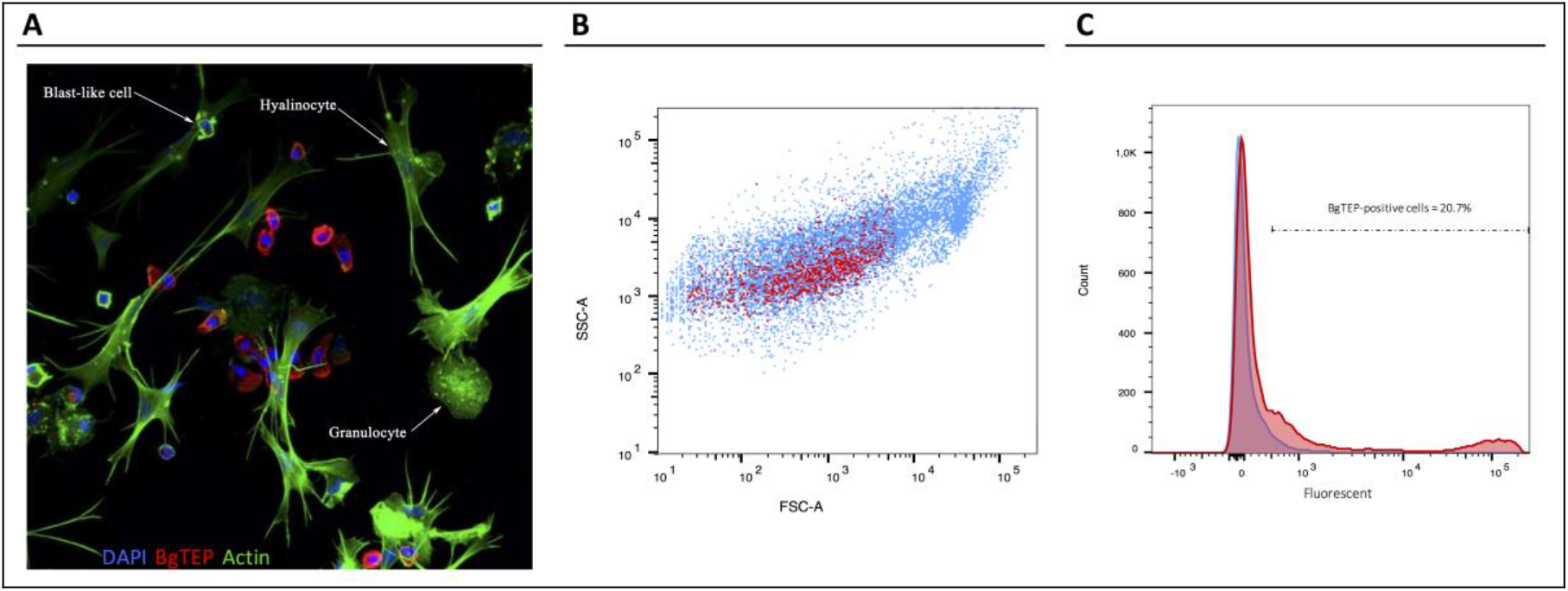
Cell-type expression of BgTEP. A. Immunolocalisation of BgTEP in hemocyte populations by confocal microscopy. Three major cell types are present in Biomphalaria glabrata hemolymph; the hyalinocytes with cytoplasmic projections, the granulocytes with granules into the cytoplasm and the blast-like cells, the smallest cells. Detection of intracellular BgTEP was performed by immunolocalisation using the antibody anti-BgTEP-PEP. Alexa-488 phalloidin was used to visualise actin filaments (green) and DAPIfor nuclear staining. The red labelling corresponds to BgTEP detection. B. Flow cytometry profile of BgTEP-positive hemocytes in hemolymph. The total hemocytes are shown according to their size FSC and granularity SSC. The red dots correspond to BgTEP-positive hemocytes. C. Flow cytometry quantification of BgTEP-positive cells present in hemolymph. Negative control was performed using the conjugated secondary antibody alone (blue). A fluorescent cut-off was determined to count BgTEP-positive cells.

### Potential role of BgTEP-positive hemocytes in phagocytosis and/or encapsulation processes

As some immune cells express BgTEP, we focused on its phagocytosis and encapsulation role. A phagocytosis experiment was performed using a yeast strain of *S. cerevisiae* conjugated to the Alexa Fluor 488 dye. The yeasts were injected *in vivo* in snails, and after 3h the plasma was recovered to observe yeast phagocytosis by confocal microscopy (Figure 8.1). Interestingly, yeast phagocytosis was observed but only in immune cells that do not express BgTEP. Among immune cells, we did not observe phagocytosis in granulocytes and blast-like cells. The same approach was applied using *E. coli* and *S. aureus* bacteria, and no phagocytosis was observed in blast-like cells, for those that expressed the BgTEP and for those that didn’t (data not shown). In conclusion, hemocytes that express BgTEP are not directly involved in the microbe phagocytosis but must instead secret an opsonin factor capable of binding to the intruder surface and of facilitating its elimination in cooperation with other immune cell sub-types.

**Figure 8:**
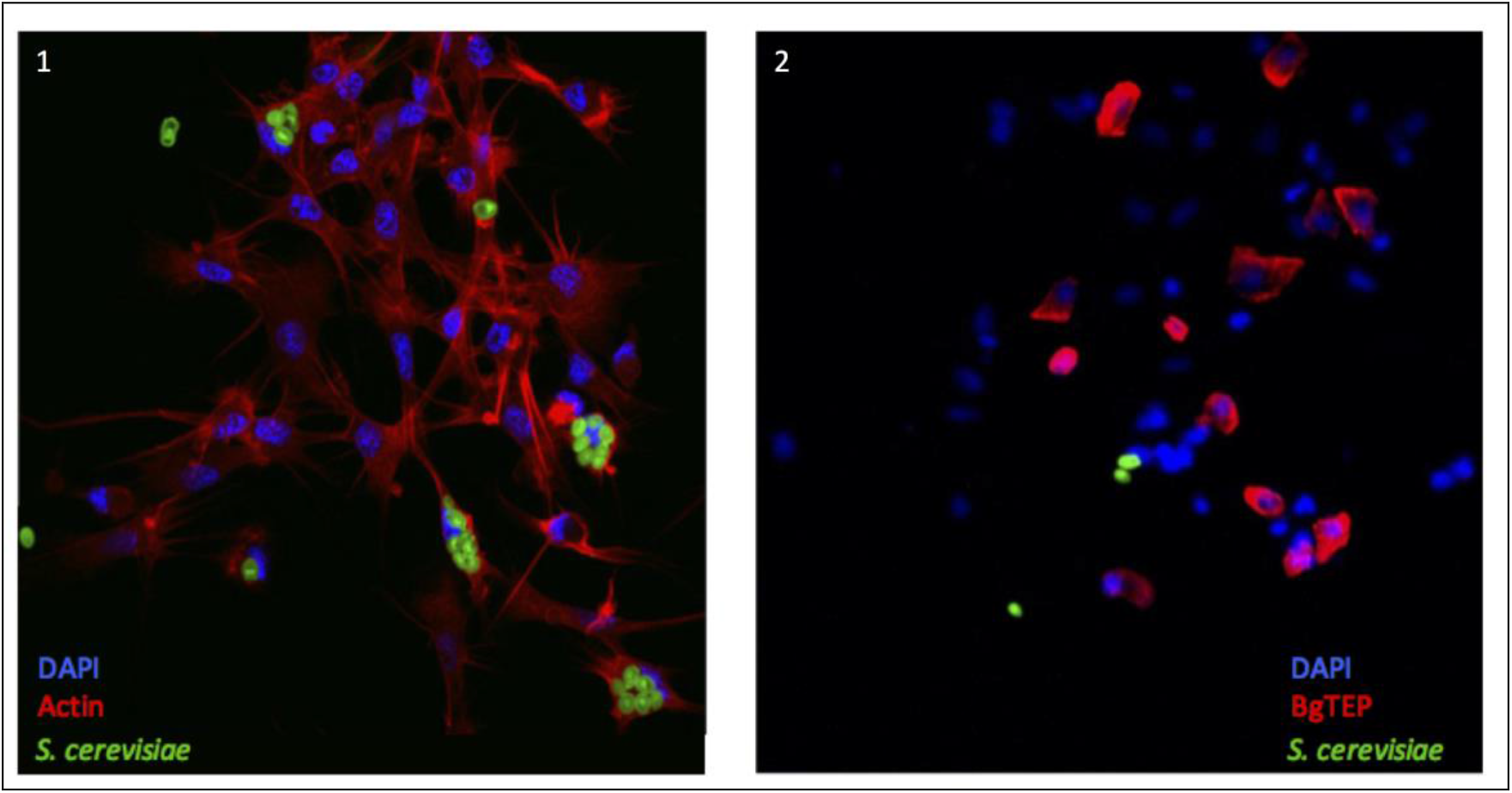
BgTEP positive cells do not phagocyte. Phagocytosis of yeasts in hemocytes observed by confocal microscopy. On picture 1, Alexa-594 phalloidin was used to label actin of all hemocytes (red). Phagocytosis assay was monitored using green fluorescent yeasts. On picture 2, BgTEP positive cells were detected by immunolocalisation using the antibody anti-BgTEP-PEP and an Alexa Fluor 594 dye conjugated to the secondary antibody (red).

Finally, the role of BgTEP in parasite encapsulation was assessed through *in situ* immunocytochemistry localisation. Following infection of *B. glabrata* by an incompatible *S. mansoni* parasite, the snail mounted a cellular immune response resulting in the encapsulation of the parasite by hemocytes. Histological sections of *S. mansoni*-infected snails were performed after 24h post-exposure. Using the anti-BgTEP-PEP antibody (Figure 8B), we observed a diffuse labelling around the parasite into the capsule. This suggests that BgTEP gets in contact with the parasite, which is consistent with the *in vitro* binding demonstration of BgTEP to primary sporocyst surface (Figure 6B). We also observed numerous BgTEP-positive hemocytes recruiting to the site of encapsulation, as well as in the vicinity of the capsule and inside the capsule (data not shown). A less intense labelling is further observed in head-foot cell wall, which is in line with the BgTEP expression measured by QRT-PCR in this tissue. These results support the hypothesis that BgTEP plays a sensing role, which could in turn promote the recruitment of other hemocyte subtypes required to mount an efficient phagocytosis or encapsulation immune response.

## Discussion

BgTEP characterisation started 25 years ago with a first report of a proteinase inhibitory activity in the plasma of *Biomphalaria glabrata* (Bender et al., 1992). In the current study, we describe an alpha macroglobulin with trypsin inhibitory activity that is sensitive to methylamine treatment (Bender and Bayne, 1996;Fryer et al., 1996). We succeeded in purifying the protein responsible for these activities and obtained the first 18 amino acids by Edman degradation sequencing. This sequence corresponded to the N-terminal part, excluding its signal peptide of BgTEP, defined in 2010 as a third actor in immune complex between snail plasmatic lectin proteins, the FREP and parasite mucins, the SmPoMuc (Moné et al., 2010). A detailed biochemical characterisation showed that this antiprotease is structured as a tetramer, and undergoes a conformational change after protease cleavage leading to the activation of the thioester bond. The foreign protease is then sequestered and cannot react with bulky substrates (Bender and Bayne, 1996).

The BgTEP protein sequence was characterised and displayed all the defining features of invertebrate TEP proteins with 8 MG domains, nested insertion of a CUB domain, the canonical TED domain, several putative N-glycosylation sites, and the C-terminal signature composed of six cysteine residues (Figure 1).

Herein we explore the molecular involvement of BgTEP in the innate immune response of the freshwater snail *Biomphalaria glabrata* against a large panel of intruders. To elucidate the nature of BgTEP within the context of the TEP superfamily, we performed a phylogenetic analysis based on full-length amino acid sequences that segregated the three main groups; complement factors, A_2_M and TEP/CD109 (Figure 2). BgTEP clearly clusters within the iTEP/CD109 group, and does not cluster with the A_2_M group, despite the previous report of BgTEP proteinase inhibitory activity (Bender and Bayne, 1996). Interestingly, a subgroup is formed with both iTEPs and CD109 from mollusks, which clusters more distantly from iTEP and CD109 proteins of all other phyla. This may be related to the primary sequence similarities between mollusk iTEPs, or may be due to a functional specificity. Among this mollusk subgroup, many proteins are predicted from automatic genome annotations, which means that they were not previously characterised, with the exception of clam TEP (Zhang et al., 2007) and bobtail squid CD109 (Yazzie et al., 2015). A unique sequence from this subgroup carries a predicted C-terminal transmembrane domain while most of them were automatically annotated as CD109 proteins. A deeper characterisation and re-annotation of each of these molecules deserves further investigation.

We then investigated the intron-exon structure of the BgTEP gene using the recently published B. *glabrata* genome (Adema et al., 2017). We identified 37 exons distributed on 10 different scaffolds from the genome assembly (BglaB1 assembly) (Figure 1). Such organisation is consistent with the one of human TEP gene families like human CD109, A_2_M and C3 complement factor genes that respectively encompass 33, 36 and 41 exons (Prosper, 2011) and the TEP of the invertebrate *Chlamys farreri* with 40 exons. Moreover, the phylogenetic analysis of the TEP superfamily reflects a mollusk TEP group that is close to vertebrate TEP and CD109 suggests a lower evolutionary divergence than with other TEPs (Zhang et al., 2009). Indeed, the BgTEP gene is considerably different from the AgTEP1 gene that is composed of only 11 exons (Personal communication from Vectorbase), thus suggesting a strikingly different evolutionary story between snail and mosquito TEPs. Despite these differences, it is not clear whether phylogenetic proximity and genomic organisation are linked to TEP function as TEP activity is more conditioned by its quaternary structure than by its primary sequence (Williams and Baxter, 2014). A structural protein prediction and alignment reveals a very close conformation between AgTEP1 and BgTEP. This result reflects a potential common function between those two complement-like components which otherwise display a low primary sequence similarity. AgTEP1, which is the most studied and the only crystallised invertebrate TEP, was shown to opsonise Gram-negative and -positive bacteria, and to promote their phagocytosis (Levashina et al., 2001). AgTEP1 was also shown to target Plasmodium parasites for lysis through a hemocyte encapsulation process (Blandin and Levashina, 2004).

The high expression level of BgTEP transcripts in snail hemocytes, the specialised circulating immune cells in *Biomphalaria* snails, correlates with results obtained for *Anopheles gambiae* (Levashina et al., 2001), thereby emphasising its potential immune function. However, high expression levels were also observed in other tissues including ovotestis. Ovotestis is the centre of production for eggs and spermatozoids, and is of paramount importance for putative immune molecule transmission to progeny. In a previous study, *B. glabrata* was shown to invest in its offspring’s protection (Baron et al., 2013) and several immune factors including BgTEP (called A_2_M when published) were recovered by proteomic analysis in egg masses (Hathaway et al., 2010). Production of BgTEP transcripts in ovotestis is thus highly relevant with this potential transfer of protection to eggs and progeny. Moreover, a recent study showed in Anopheles that during spermatogenesis, AgTEP1 binds to and removes damaged cells, increasing fertility rates (Pompon and Levashina, 2015).

In naïve snails, BgTEP is constitutively secreted in the hemolymph, and expressed at high levels in circulating immune cells. Western blot on plasma, using the anti-BgTEP-PEP antibody, revealed the presence of full length, as well as several processed forms, of BgTEP. This suggests that BgTEP undergoes a cleavage by an undefined plasmatic factor, similar in the case of AgTEP1 (Levashina et al., 2001; Shokal et al., 2017). In *A. gambiae*, the AgTEP1 is found in full-length and in a processed form called TEP-cut, which allows for a complex pattern of TEP that is ready to respond to a pathogen attack (Levashina et al., 2001). In hemolymph, the AgTEP1 is maintained by a complex of two proteins APL1 and LRIM1 to stabilise the processed form and to avoid the unspecific binding of the thioester domain to non-relevant substrates (Levashina et al., 2001;Shokal et al., 2017). In vertebrates, the complement component pathway displays a major role in the innate immune system. The complement component C3 activation is finely regulated by a series of proteolytic cleavages leading to the formation of different fragments of C3 such as C3a, C3b, iC3b, and C3dg. All proteolytic fragments, such as,the small complement fragment C3a, mediate chemotaxis and local inflammation. C3b acts as an opsonin by enhancing cellular phagocytosis by binding to the pathogen’s surface. The C3b derived fragment, iC3b and C3dg can bind to the pathogen and promote its uptake (Hamad et al., 2010;Feng et al., 2015;Foley et al., 2015).

Interestingly, an immune labelling on hemocytes revealed that only a subset of blast-like cells is positive for BgTEP (Figure 7A). This observation possibly suggests that more hemocyte subtypes or a differential maturation exist in the *B. glabrata* hemolymph as compared to the ones previously estimated solely on the basis of cell morphology analysis. Further investigations are needed to confirm this observation through a characterisation of functions for each hemocyte type.

Through a targeted interactome approach, we observed that BgTEP is retrieved bound to all intruders tested in the present study, in its full length as well as in processed forms (Figure 6). This suggests that the full lengh of BgTEP bound on the intruders surface can be cleaved by a proteolytic cascade or that processed forms can bind directly. These results are consistent with the previously observed binding of AgTEP1 to bacteria that was shown to occur in both thioester-dependent and thioester-independent scenarios (Levashina et al., 2001 Cell). This could also indicate that BgTEP is probably able to bind intruder surfaces directly or indirectly associated with other immune relevant partners. After 3 hours of interaction, fewer full length and 30 kDa processed forms of BgTEP were recovered in forms bound to the surface of *E. coli, S. cerevisiae* and *S. mansoni*, indicating a time-dependent processing compared to a shorter treatment (Figure 6). New forms of BgTEP appeared after 3 hours of incubation with yeast and bacteria, which could result from the processing of already bound full-length protein (Figure 6A). Another striking result is the difference observed for the binding of BgTEP between miracidia and sporocysts, which are two successive developmental stages of *S. mansoni* parasite (Figure 6B). Interestingly, more processed forms of BgTEP were recovered bound to sporocysts than to miracidia, and no bands were detected with miracidia after 3 hours, suggesting a higher specificity of BgTEP for sporocyste stage than for miracidia. Such a result is not surprising as a sporocyst is the result of miracidia transformation which consists mainly of the loss of epidermal ciliated plates and tegument renewal that occurs in the early hours after infection. Several proteomic and glycomic studies have highlighted differences from one developmental stage to another (Hokke et al., 2007 Trends in Parasitol; Peterson et al., 2009 Int J of Parasitol). These results would suggest a subtle ability of the snail immune machinery to distinguish between various developmental stages of the parasite.

Collectively, these results clearly indicate that BgTEP can be associated with the surface of live intruders and could be differentially processed depending on the intruder type. For the first time, we also approached the dynamic of immune complex formation with a selective processing of bound TEP between intruders. So even if intruders were sensed by this complement-like factor (Tetreau et al., 2017), other maturation factors may be involved to induce an appropriate immune response. Nevertheless, the binding mechanisms are still unclear and need to be deeply characterised.

As BgTEP can interact with several intruders, we investigated the relative expression of BgTEP transcript by quantitative RT-PCR following various immune challenges (Figure 5). BgTEP expression is modulated, regardless of the nature of the intruders used for the stimulation step. *E. coli* and *S. cerevisiae* challenges decreased its expression, while *M. luteus* and *S. mansoni* challenges up regulated it, as previously detailed in the transcriptomic analysis of the snail immune response after bacterial and fungal infections (Deleury et al., 2012). Interestingly, *S. mansoni* is the only one that induced a constant increase from 6 h to 24 h of BgTEP transcript expression suggesting a role of first importance in the interaction between *S. mansoni* and *Biomphalaria glabrata*, which supports the first identification of BgTEP in a host-parasite immune complex (Moné et al., 2010). Although we observed many BgTEP-positive hemocytes converging towards the encapsulated parasite and surrounding the hemocyte capsule, a typical associated immune response with an incompatible strain of *S. mansoni*. This observation suggests the participation of BgTEP-positive hemocytes in the recruitment of capsule-forming hemocytes on the site of infection, potentially via a putative α-2-macroglobulin receptor on their membrane (Coustau et al., 2015;Paul et al., 2016;Pila et al., 2017). Further, hemocytes converging to the site of infection may also indicate a chemotaxis property of BgTEP due to a cleavage of bound TEP into small anaphylatoxin-like fragments.

This study provides new insights about the potential immune function of BgTEP. We demonstrate that its constitutive production by hemocytes must be modulated by immune challenges, and that the full protein and its proteolytic fragments are able to bind the surface of different intruders before and after specific cleavage maturing processes. Even though the precise binding mechanism needs further characterisation, our results suggest that BgTEP displayed an immune role by targeting intruder surface.

In this work, we report the first characterisation of an iTEP displaying a dual-role, whose existence was previously argued (Williams and Baxter, 2014). As described before, BgTEP acts as an antiprotease (Bender and Bayne, 1996), but in this study we demonstrate that BgTEP can also bind to different intruders, including the *S. mansoni* parasite, and could participate in their elimination.

In conclusion, a more precise functional characterisation is necessary to decipher the key role of the BgTEP and its action dynamics during the immune response of the snail. To that end, a loss of gene function by CRISPR/cas9 technology or RNAi would be considered. Also, the nature and function of proteolytic products of BgTEP remain unknown and must be explored to elucidate host-pathogen interaction. Indeed, some pathogens circumvent the host immune response by blocking or miscleaving complement components (Jusko et al., 2014;Johnson et al., 2015;Luo et al., 2018).

## Acknowledgements and funding information

We thank Dr. Chaparro Cristian and Duval Jérôme for time spent reading and improving our manuscript. The authors would like to thank Damien Lassalle for his help for snail breeding. We thank the referee for his relevant and constructive comments that were very helpful to improve the manuscript.

This work was supported by the French National Agency for Research (ANR) through a project Invimory grant [ANR-13-JSV7-0009] to BG.

